# *Mdx* mice dosed with antisense to CD49d and dystrophin exon skip morpholino; reduced force loss and affected gene expression pathways support ATL1102 combination therapy in DMD patients

**DOI:** 10.1101/2024.11.23.625028

**Authors:** A. S. Padhye, L. Kiriaev, C. Tiong, L.J Gearing, T Wilson, P. J. Houweling, G. Tachas

## Abstract

**Background:** VLA-4 is a disease progression biomarker in patients with Duchenne Muscular Dystrophy (DMD) including those treated with corticosteroids. ATL1102 an antisense oligonucleotide (ASO) inhibitor of human CD49d chain of adhesion molecule VLA-4 shows promising phase-2 results in non-ambulant DMD stabilizing upper limb function (PUL2.0), grip strength, and MRI muscle fat fraction, compared to worsening with corticosteroids.

**Methods:** ASO to mouse CD49d (ISIS348574) was used in combination with a known mdx dystrophin morpholino exon-23 skipping restoration drug (PMO), dose-time chosen for low dystrophin restoration. *Mdx* mice were treated once weekly for 8 weeks with ASO, control mismatch oligonucleotide, saline, or PMO alone for 4 weeks, and PMO in combination with ASO or mismatch.

**Results:** ASO+PMO demonstrated increased specific maximum force and eccentric muscle force retention in the exterior digitorum longus (EDL) muscle after 1-7 muscle eccentric contractions relative to saline. Area-under-curve force remaining after 8-10 contractions was higher for ASO+PMO versus PMO monotherapy and ASO monotherapy, PMO monotherapy versus saline, and the ASO monotherapy versus saline and mismatch.

RNA-sequencing of quadriceps muscle evaluated gene expression changes, with false discovery rate (FDR) adjusted p-values up to <0.1. ASO monotherapy at FDR<0.05 showed 2 unique genes, *Gm2a* in immune neutrophil function and *Trdn* with *Ryr2* muscle calcium channel function, versus *mdx* saline and none with MM. The lowest FDR PMO gene was another calcium channel *Cacna1s*, with PMO+MM FDR<0.05. ASO+PMO treatment at FDR<0.05 modulated 55 genes, 53 unique to the combination and 2* also in ASO FDR<0.06. Affected genes were involved in immune response (*Btla*\*, *Ppml1*, *Tnfsm13*), lipolysis (*Fabp4*\*, *G0s2*), fibrosis (*Igfbp-7, Calu*), muscle cell (*Asb15*) and muscle stem cell function (*Adam10, Mt-Tp, Myom1*).

**Conclusion:** ASO+PMO EDL muscle function protection in *mdx* mice, and gene expression pathways affected, support the development of ATL1102 in combination therapy with conditionally approved morpholino dystrophin restoration drugs in DMD patients.

## Introduction

Duchenne Muscular Dystrophy (DMD) is a neuromuscular X-linked disorder that affects approximately 1 in 5000 live male births worldwide [1]. The primary cause of DMD is a mutation in the dystrophin gene, which results in the loss or absence of functional dystrophin in muscle and secondary immune-mediated inflammatory damage, fibrosis, fatty infiltration and loss of stem cells in muscle. DMD is the most common genetic disorder in boys and causes progressive muscle weakness in the first decade of life, leading to loss of ambulation in the second decade, with non-ambulant boys needing ventilation in the second decade, and death often resulting due to cardiorespiratory failure in the third decade [2].

Corticosteroid (CS) therapy targeting the inflammatory damage in DMD is the standard of care in DMD but delays loss of ambulation by only 1 – 3 years in boys to 11 – 13 years of age, and treatment is associated with a myriad of adverse effects [3]. Dystrophin restoration drugs [4], such as the morpholino antisense exon skipping drugs eteplirsen [5], viltepso, golodirsen [6], and casimersen, produce small amounts of dystrophin and are conditionally approved in the US and used with CS in patients with certain dystrophin mutations [7].

Ataluren, a stop codon readthrough dystrophin restoration drug, similarly produces small amounts of dystrophin, and has purported benefits on respiration and ambulation on top of CS but registration in the European Union remains under re-evaluation due to lack of efficacy on ambulation endpoints [7–9]. There is a need for more effective and safe treatments for ambulant boys with DMD targeting the primary, secondary and other causes of disease to delay disease progression and stabilize limb function including in combination with the first-generation exon skipping drugs.

Loss of functional dystrophin in myofibers in DMD increases susceptibility to calcium contraction-induced membrane damage and ultimately leads to chronic inflammation. Inflammation involves activation of the innate immune system, with upregulation of pro-inflammatory cytokines [10] and extracellular osteopontin expression in muscle fibers and inflammatory infiltrates, followed by the activation of the adaptive immune system.

Increased expression of the very late antigen 4 (VLA-4) adhesion molecule on a greater number of circulating T cells in the adaptive immune system is associated with more severe and progressive disease related to uptake of such cells in muscles [11,12]. Patients with DMD expressing high levels of the VLA-4 ligand osteopontin in muscle are also not as responsive to CS, and osteopontin plays a role in fibrosis in in ambulant patients with DMD [13,14].

ATL1102 is a second-generation antisense oligonucleotide (ASO) to human CD49d that reduces circulating CD49d+ T cells and in a small study, was shown to stabilize limb function, strength, and muscle structure in non-ambulant DMD subjects compared to worsening with CS. ATL1102 has been shown to stabilize the performance of upper limb function PUL2.0 score and MyoPinch and MyoGrip strength measurements compared to losses with CS, reduces MRI detected fat fraction, and increases lean muscle mass compared to worsening reported with CS [15–17]. In the *mdx* mouse model of DMD, symptomatic 9-week old mice treated for 6 weeks with ASO to mouse CD49d RNA (ISIS348574), the murine equivalent to ATL1102, have reduced CD49d mRNA expression in quadriceps muscle and reduced *tibialis anterior* skeletal muscle function loss post eccentric muscle damage [17].

The current study is the first to investigate the potential of combining the immune-modulating effects of murine ATL1102 (ISIS348574) to CD49d with a known dystrophin morpholino exon 23 skipping restoration drug (PMO), at a dose and time previously shown to produce dystrophin at low levels in *mdx* mouse muscles, to parallel low level dystrophin production in DMD subjects with first generation exon skipping drugs. The PMO when previously used in *mdx* to produce high 8.6% dystrophin levels in the exterior digitorum longus (EDL) [18] and 10-20% dystrophin in *tibialis anterior* [19], reduces loss of function in *mdx* vs saline, but this is at higher dystrophin levels than produced with exon skipping drugs in DMD patients. The present study showed the ASO+PMO combination benefits on the EDL muscle compared saline. The multiple pathways affected in quadriceps muscle are reported which indicate benefits in immune, fibrosis, lipolysis and muscle cell and stem cell pathways, and support future trials of ATL1102 with PMO dystrophin restoration drugs in patients with DMD.

## Methods

### Oligonucleotides

The ASO targeting mouse CD49d (ISIS348574), a second-generation 2′-*O*-methoxyethylribose (MOE) gapmer fully phosphorothioate oligonucleotide, and negative control mismatch (MM) oligonucleotide with the same second-generation MOE gapmer chemistry and nucleotides in a different order to ISIS348574 (16) were manufactured by Integrated DNA Technologies. The PMO targeting dystrophin pre-mRNA (+7 18; 5-GGCCAAACCTCGGCTTACCTGAAAT-3) exon 23 skipping restoration drug was manufactured by Gene Tools, LLC [20]. These oligonucleotides were used at the same doses as per published studies in *mdx* mice. The ASO to mouse CD49d (ISIS348574) previously used at 20mg/kg s.c once weekly doses in 9-week old male B10.*mdx* (*mdx*) mice for 6 weeks, assessed soon after the last dose, reduces target CD49d in quadriceps muscle by 40%, reduces CD4+ and CD8+ T cells as a percent of all live cells in the spleen (30% and 20% respectively) and improves by ∼25% tibialis anterior force remaining *in situ* following eccentric muscle contraction versus control *mdx* mice [17].

The PMO targeting dystrophin pre-mRNA (+7 18; 5-GGCCAAACCTCGGCTTACCTGAAAT-3) exon 23 skipping restoration drug used at 50mg/kg i.v once weekly doses for 3 weeks and followed for over 4 weeks post dosing produces dystrophin in *mdx* mice at similar low levels seen with PMO targeting human dystrophin exon splicing restoration drugs eteplirsen and golodirsen and viltolarsen in boys with DMD [20]; generating at 17 days post dosing ∼4% of WT dystrophin levels in quadriceps, with diminution by day 31 to ∼1.5% dystrophin, which remains distinguishable from baseline levels, as shown in Novak *et al* (2017, supplemental Fig. 2.B) [20].

### Animals and ethics approval

All animal procedures in this study were conducted in accordance with guidelines following approval by the Animal Ethics Committee of the Murdoch Children’s Research Institute (MCRI) animal care and ethics committee (ACEC), approval number ID# A899.

Male wild type (WT) C57BL.10 control and B10.*mdx* (*mdx*) age matched mice were purchased from Jackson laboratories at 5 weeks of age and acclimatised to the MCRI facility for a total of 4 weeks. Animals were housed in a specific-pathogen-free environment at a constant ambient temperature of 22°C and 50% humidity on a 12 h light-dark cycle, with *ad libitum* access to standard chow food and water.

### Animal study groups and sample collection

Male WT C57BL10 mice and male B10.*mdx* (*mdx*) mice 9 weeks of age were treated for 8 weeks as described below and as shown in Fig. 1; once weekly saline (WT group 1 and *mdx* group 2), ASO to mouse CD49d (ISIS348574) (group 3), negative control mismatch (MM) oligonucleotide (group 4), PMO drug alone for 4 weeks and saline for another 4 weeks (group 5), ASO+PMO (group 6) or PMO+MM (group 7).

**Fig. 1.**
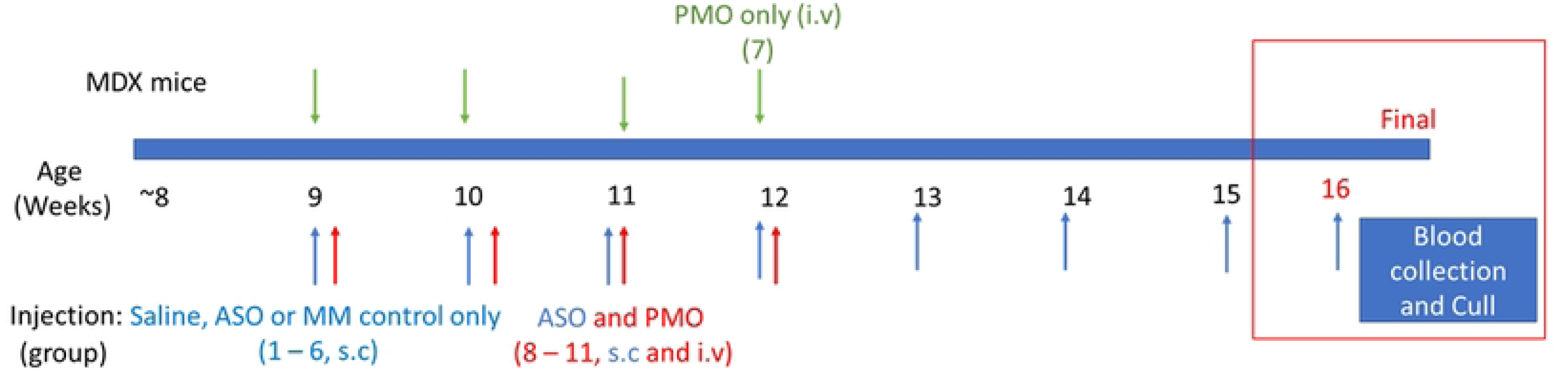
Mdx mice treatment with saline, MM control, ASO, PMO monotherapy and MM+PMO, and ASO+PMO combinations as subcutaneous (s.c), and intravenous (i.v) injections.

A total of 84 mice were used across the 7 groups (n=12 per group) and mice were treated for 8 weeks and samples collected as follows:

1. WT C57BL10 mice dosed with saline s.c once weekly for 8 weeks (WT Control), with sample collection 2 days past the last dose.
2. *Mdx* mice B10.*mdx* dosed with saline s.c once weekly for 8 weeks (Saline), with sample collection 2 days past the last dose; *mdx* mice have the WT background phenotype.
3. *Mdx* mice dosed with ISIS348574 antisense to CD49d (ASO) (20mg/kg) s.c once weekly for 8 weeks, with sample collection 2 days past the last dose.
4. *Mdx* mice dosed with negative control mismatch (MM) oligonucleotide (20mg/kg) s.c once weekly for 8 weeks, with sample collection 2 days past the last dose.
5. *Mdx* mice dosed with dystrophin morpholino exon 23 skipping restoration drug (PMO) 50mg/kg i.v into the tail vein once weekly for 4 weeks, followed by saline s.c once weekly for 4 weeks, with sample collection 31 days past the last dose of PMO, and 2 days past the last dose of saline.
6. *Mdx* mice dosed with ASO 20mg/kg s.c once weekly for 8 weeks, and PMO 50mg/kg i.v for the first 4 weeks, with sample collection 31 days past the last dose of PMO, and 2 days past the last dose of ASO.
7. *Mdx* mice dosed with MM 20mg/kg s.c once weekly for 8 weeks, and PMO 50mg/kg i.v once weekly for the first 4 weeks, with sample collection 31 days past the last dose of PMO, and 2 days past the last dose of MM.

After treatment samples were collected, mice were anaesthetised using inhaled isoflurane and then euthanised by cervical dislocation once animals were no longer breathing. Muscle samples were surgically removed, EDL for the muscle functional study, and quadriceps for dystrophin protein, and fibre immunofluorescence and RNA sequencing studies using RNA-Seq. Not all 12 animals per group were utilized for each experimental measure, with specific subsets selected at random for individual analyses based on the experimental design and the requirements of each procedure.

### Muscle functional study

For the muscle functional study, *ex vivo* EDL muscle physiology was performed in mice in each group following Kiriaev et al 2018 with minor modifications [21]. The hindlimb fast-twitch EDL muscle was dissected in carbogenated Krebs solution, and one end was tied to a force transducer/linear tissue puller using the tendons and the other end secured to a fixed pole.

An initial supramaximal stimulus was given at 1 ms pulse width, 125 Hz for 1s duration, and force produced for muscles recorded, and the maximum force output of the EDL muscle at optimal/resting length recorded. Force-frequency curves were generated by trains of stimuli given 30s apart at different frequencies, including 2, 15, 25, 37.5, 50, 75, 100, 125, 150 Hz. A sigmoid curve relating the muscle force to the stimulation frequency was fitted by linear regression to this data using Graph Pad Prism as done in Kiriaev et al 2018 [21]. Following Hayes and Williams 1998 [22], specific force was calculated according to the equation Cross sectional area (CSA) = muscle mass/(Lo × D), where Lo is the optimal length and D is the density of skeletal muscle (1.06 g/cm3).

Following a 5-minute rest, 10 eccentric (lengthening) muscle contractions (EC) were conducted where the muscle was stretched 10% from the optimal fibre length (Lo). At t = 0 s, the muscle was stimulated via supramaximal pulses of 1 ms duration (width) and 125 Hz frequency. At t = 0.9 s, after maximal isometric force was attained, each muscle was stretched 10% longer than its optimal length. The muscle was then held at this length for 2 s before returning to optimal length. Electrical stimulus was stopped at t = 5 s. This procedure was performed 10 times at intervals of 3 min. The force measured at each EC was expressed as a percentage of the force produced during the first (initial) contraction.

### Statistical analysis of EDL functional data

Analyses of Maximum specific force were performed in GraphPad Prism (V9, Graphpad Software Inc.) using a One-way ANOVA with Tukey’s (Fig. 2 A) with a planned hypothesis on assessing the combination of ASO+PMO and MM+PMO combination versus salines shown in Fig.2A Analyses of force remaining after 5 EC were performed across the groups using a One-way ANOVA with Tukey’s and Dunnett’s (Fig. 2B). Data was also analysed over the 10 EC using a Two-way ANOVA with Tukey’s focussed on the planned hypothesis of ASO+PMO and MM+PMO combinations versus saline (Fig. 2B) and data presented as bar graphs with means ± SD error bars for each group. Data over the 10 EC were also presented as a smoothed line with the calculated group mean at individual points (Fig. 2D) with the Area Under the Curve (AUC) presented as a bar graph with means ± SD error bars for each group, as analysed using a One-way ANOVA with Tukey’s correction after 8 and 10 EC respectively (Fig. 2F and 2G).

**Fig. 2.**
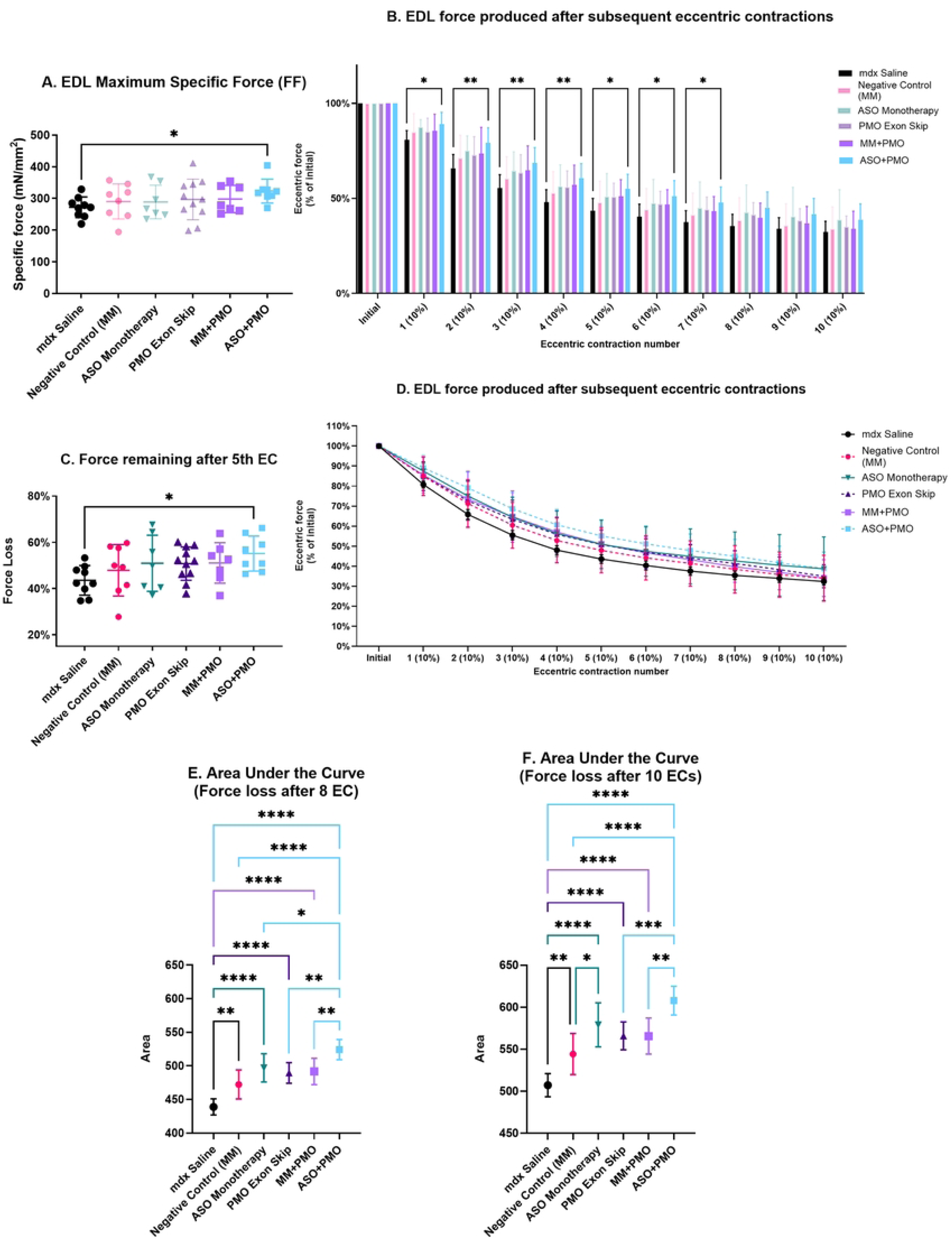
*Mdx* mice treated with ASO to CD49d in combination with PMO resulted in the EDL maximal specific force increase and decreased force loss after eccentric lengthening contractions. (A) An initial supramaximal stimulus was given at 125Hz (1ms pulses) for 1s and force produced recorded as the maximum force output of the EDL fast-twitch muscle relative to the Cross Sectional Area at Lo. Mean 18.8% significant differences were observed between saline treated *mdx* mice (272mN/mm2) and the ASO+PMO combination (323.3 mN/mm2) (n=7-11, One-way ANOVA with Tukey’s test, p = * <0.05) (B) A series of 10 eccentric contractions (EC) were performed on each EDL where the contracted muscle was stretched by 10% and held for 2s before returning to L. The force output relative to baseline is shown as a bar graph % of initial force produced. The figure shows the differences of the ASO+PMO combination after the 1st, 2nd, 3rd, 4th 5th, 6th, 7th EC were significant differences versus saline p = * <0.05 and p=**<0.01. (Repeat measure Two-way ANOVA with Tukey’s test in a planned comparison vs MM+PMO and saline). After the 1st EC, the ASO+PMO combination showed reduced force loss (10.8%) compared to the saline *mdx* mice (19.2%). (C) After the 5th EC, the ASO+PMO combination showed ∼20% reduced force loss (44.9%) compared to the saline *mdx* mice (56.4%) (One-way ANOVA with Dunnett’s test, p = * <0.05). (D) The force output relative to baseline after each of the 10 EC shown for each groups is shown as a smoothed line graph % of the initial force produced to show the slope of early and late EC changes. (F, G) An area under the curve (AUC) calculation was performed on (D) using Graphpad Prism ®. The cumulative integrated effect across all 10 contractions (Fig. 2G) shows the various significant reductions in AUC muscle force losses from baseline. The ASO+PMO combination AUC were significantly higher versus all other treatments with trends vs ASO (p=0.076) after 10 contractions, that were significantly different earlier after 8 contractions Fig. 2F (p=0.0045). The ASO monotherapy was greater than saline and MM, and the PMO was greater than the saline. (One-way ANOVA with Tukey’s correction, p = * <0.05, ** <0.01, *** <0.001), ****<0.0001) eccentric contractions (p=0.0045) Fig. 2F. The ASO monotherapy showed a reduction in AUC force loss compared to the MM and saline treated *mdx* mice, and the PMO monotherapy showed a reduction in AUC force loss compared to saline treated *mdx* mice (Fig. 2G). Looking at AUC after 4 eccentric contractions there was no significant differences between the MM and saline treated *mdx* mice.

### Quadriceps dystrophin protein expression and fibre dystrophin intensity quantitation

Dystrophin protein expression data was determined by western blot using the ProteinSimple capillary Western Immunoassay (Wes) and monoclonal dystrophin antibody (ab154168, 1/1000 dilution) as previously described [23,24]. Snap-frozen quadriceps muscles (n = 6 / group) were homogenized in 2% SDS lysis buffer and assessed for total protein concentration using Direct Detect (ThermoFisher Scientific). To verify total protein detection and adjust for loading, the total protein detection module (DM-TP01; ProteinSimple, biotechne) was used, ensuring precise and quantitative analysis of protein expression levels.

The Wes was performed (SM-W008; ProteinSimple, biotechne) according to the manufacturer’s instructions using the 66-440 kDa (pre-filled plate) separation module. The rabbit monoclonal anti-dystrophin antibody (no. ab154168; Abcam; dilution 1/1,000) was used for dystrophin detection. An anti-rabbit secondary antibody was used (DM-001; Anti-rabbit detection module). The lysed tissue samples were subsequently diluted to 0.25 μg/μl, with 5 μl being loaded per cartridge well, resulting in a loading amount of 1.25 μg per well/capillary. To control for differences in signal between experiments, WT C57BL6 muscle was used to generate a 6-point standard curve (1.25μg to 0.008μg total protein loaded/well) and compared to treated *mdx* mouse muscles (1.25μg total protein). The final dystrophin values were normalized against WT and corrected for total protein.

Fibre Dystrophin fluorescence intensity was assessed using 10-12μm cryosection of the quadriceps muscle stained with polyclonal antibody to dystrophin ab15277 (1:200) as per Novak et al [20,24]. Sections were stained with rabbit anti-dystrophin antibody (no. ab15277; Abcam; dilution 1:200) and 4’,6-diamidino-2-phenylindole (DAPI) for nuclear staining (H3570; Invitrogen; dilution 1:200), diluted in phosphate-buffered saline (PBS), and incubated with the sections for 2 hours at room temperature. Sections were washed (3×) in PBS for 3 minutes each. Sections were then washed and probed with donkey anti-goat IgG (A32814; Invitrogen, Thermo-Fisher). These were diluted in PBS and incubated for 1 hour in a dark at room temperature. Sections were then washed in PBS, fixed in 4% paraformaldehyde (15735-90; ProSciTech) and mounted using Immu-Mount (9990402; Shandon, Thermo-Fisher).

Microscopy was performed using a widefield microscope (Axio Imager 2 system, ZEISS) with Zen microscopy software (v3.8, ZEISS). Analysis and quantification were performed using FIJI ImageJ software (v2.8). The Quadriceps samples were collected from matched regions of the same muscles by collecting cryosections (CM1900; clinical cryostat, Leica). A total of 21 quadricep muscles (n=3 per group) were sectioned at 10-12μm for 6 serial sections per muscle and stored at –80◦C for processing. For each stained section, 8 sample regions (800μm x 800μm) were taken from each quadricep muscle and averaged to produce a single data point representing that quadricep muscle.

### Quadriceps RNA isolation, quantification, library preparation and sequencing

Muscle RNA samples from WT C57BL10 mice 16 weeks of age and treated *mdx* mice at 16 weeks of age were used to evaluate gene expression changes.

Total RNA was extracted from ∼50 mg of snap frozen cryopulverised quadriceps muscle (n=6 mice per group) in 1 ml of TRIsure solution (Bioline). Following this, RNA was purified using the RNeasy Mini Kit and DNase treatment (QIAGEN) as per manufactures instructions and eluted in 30 μl of Milli-Q water. RNA integrities [RNA integrity number (RIN) value] and total RNA concentrations were measured using TapeStation (Agilent Technologies 2200).

A custom in-house RNA seq multiplex method was used similar to that previously described [25,26]. Samples were given a unique i7 index together with a Unique molecular identifier (UMI) during individual pA priming and first strand synthesis which also adds a template switch sequence to the 5’-end. Samples were then pooled into sets and amplified using P7 and an oligo which binds the template switch sequence. Final library construction was completed by tagmentation and addition of unique P5 (with i5 index) by PCR. Sequencing was performed on an Illumina NextSeq 2000 run with 61nt SR (cDNA). An 18 nucleotide nt i7 read contains the 8nt index and 10nt UMI. Samples were parsed using the i7 and i5 indexes.

### Statistical and pathway analysis of quadriceps RNA-seq data

The RNA-seq data analysis was performed using R [27] with quality control done using FastQC [28]. The read alignment was performed using the R subread package (v2.8.1) [29]. An index was built using the Ensembl Mus musculus GRCm38 primary assembly genome file and alignment performed with default settings with the majority of reads (78.4%), being assigned to a gene. Raw gene expression distributions were relatively similar across all samples. Before normalization, low expressed genes < 1 count(s) per million (cpm) were filtered out as per other studies [30], leaving 21,853 out of 55,487 total genes for differential analysis. An additional assessment was done with genes < 5 cpm filtered out as per previous published studies [31,32], and it was noted that 18,225 remained out of 55,487 total genes. Normalization factors were calculated using the Trimmed Mean of M Values (TMM) method [33].

Normalized gene expression data was analyzed in R using limma [34] and edgeR [35]. An eBayes moderated t-statistics test was used to test differential expression, comparing the *mdx* Saline mice to the treated *mdx* mice and WT Saline [36]. To correct for multiple comparisons, the Benjamini-Hochberg method was used to calculate the false discovery rate (FDR) adjusted p-values [37] and significantly expressed genes were identified with an FDR <

### 0.05 and thereafter with an FDR < 0.1

A pathway analysis (Genetrails3) was performed on June 2023, using the protein markers with an FDR <0.1, using an over-representation analysis with Benjamini and Hochberg adjusted p-value < 0.05.

## Results

### Muscle functional study: ASO to CD49d plus exon skip PMO improved muscle force and reduced eccentric contraction induced damage in *mdx* mice

Muscle physiology analyses of the EDL muscle showed the ASO plus PMO exon skip combination increased the specific maximum force vs saline (p=0.0271), in contrast to the MM+PMO group vs saline in a planned analysis of these combinations (Fig. 2A). Mean 18.8% significant differences were observed between saline treated *mdx* mice (272mN/mm2) and the ASO+PMO combination (323.3 mN/mm2) with n=7-9 in these groups, One-way ANOVA with Tukey’s test, p = <0.05).

ASO+PMO increased the eccentric muscle force remaining following 5 lengthening eccentric contractions (EC) versus saline treated *mdx* (Fig. 2C), Dunnett’s (p=0.0422), Tukey’s (p<0.1) with n=7-9 in these groups. After the 5th EC, the ASO+PMO combination (-44.9%) showed a significant 20.0% lower EDL force loss compared to the saline *mdx* mice (-56.4%). After the 5th EC, the mean force remaining was ASO+PMO (55.12%), MM+PMO (51.06%), PMO (51.05%), ASO (50.94%) and saline (43.46%).

Examining the area under the curve force remaining after up to 8 and after 10 eccentric contractions, showed the ASO+PMO combination was significantly higher than the PMO alone and MM+PMO, and also to the ASO alone, the latter significantly different after 8

### Dystrophin protein expression and fibre dystrophin intensity

Western blot was used to assess the level of dystrophin expression 31 days post the last dose of PMO in the saline, PMO and PMO+ASO treated muscles. All samples fell below the level of detection which was determined to be 1.5% of WT dystrophin levels (Fig. 3A), with ∼0.5% of dystrophin levels in PMO and PMO+ASO treated muscles (Fig. 3B). No dystrophin protein was detected in the *mdx* saline control group (Fig. 3B).

**Fig. 3.**
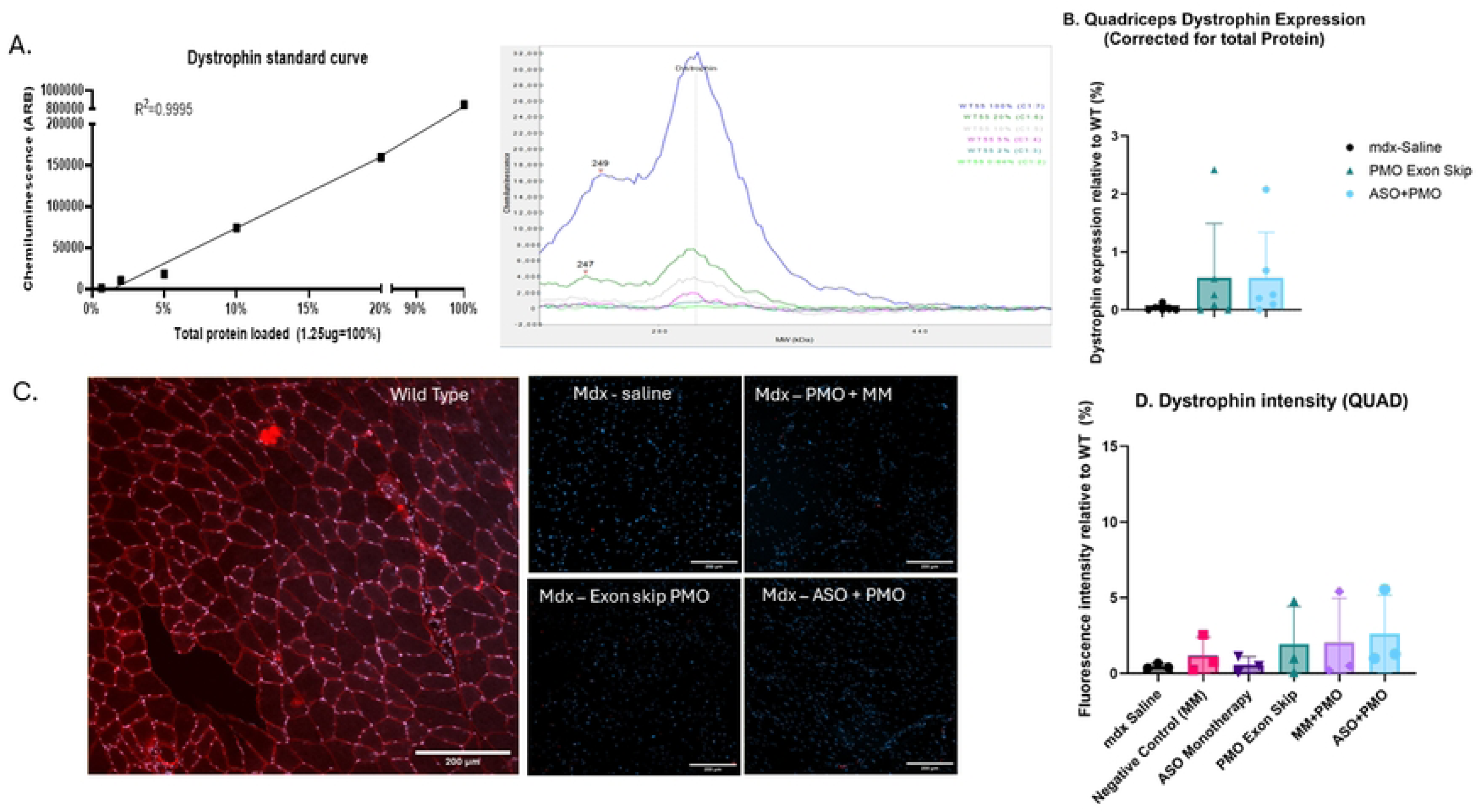
Western blot quantification of dystrophin levels in the *mdx* versus WT mice and muscle fibre dystrophin immunofluorescence intensity data. (A) Standard curve of diluted dystrophin from WT muscle with representative chromatograph from the WES ProteinSimple analysis tool. (B) Dystrophin levels in PMO monotherapy and ASO+PMO treated *mdx* mice and *mdx* saline treated controls relative to WT using ProteinSimple (WES) and (C) Representative images of cross sectional quadricep muscle dystrophin in WT, *mdx*-saline, *mdx*-PMO+MM, *mdx*-PMO and *mdx* ASO+PMO (Dystrophin (Red), DAPI (Blue). (D) Quantitation of dystrophin fluorescence intensity using IHC including PMO monotherapy ASO+PMO and MM+PMO groups relative to *mdx* saline treated mice.

Dystrophin intensity in the muscle fibres was assessed in all groups using immunofluorescence in matched quadricep tissue sections. A similarly low percentage of dystrophin fluorescence was observed in the PMO+ASO combinations and the PMO monotherapy treated mice, with less than 2.5% of WT dystrophin levels and no dystrophin fluorescence in the ASO, MM, or saline control (Fig. 3C and D).

### WT versus mdx saline quadriceps gene expression changes associated with immune cells, fibrosis, and muscle cells

A total of 21,853 transcripts were identified with 6464 genes expressed significantly different between WT versus *mdx* saline at FDR <0.05 in the 16-week-old male mice. At FDR <0.0005 there were 2562 genes differentially expressed. Various CD49d ligand immune and fibrosis genes were differentially expressed in *mdx* mice versus WT, with *mdx* showing increased expression of CD29 *Itgb1* (1.4-fold), *Spp1* (69-fold), *Fn1* (3.7-fold) and *Tgfβ1* (4.2-fold) though reduced expression of CD49d *Itga4* (0.4-fold). Increased expression of CD49d ligand *Fn1* for fibrosis and *Spp1* for calcification are previously reported in areas of muscle regeneration [38]. Genes involved in muscle physiology differentially expressed in *mdx* mice versus WT in this study included *Myostatin* (down 0.2-fold), *Dmd* (down 0.5-fold), *Igf-1* (up 1.9-fold), and *Myod1* (up 2-fold).

Two proteins recently targeted in phase 3 clinical trials in DMD patients are CTGF and HDAC1 involved in fibrosis and immune cells[39,40]. *Ctgf* was not differentially expressed and *Hdac1* was increased by 1.8-fold in the *mdx* versus WT mice.

### ASO monotherapy changes in gene expression linked to calcium handling, immune function, fibrosis, lipolysis, muscle stem cells and contraction

In the ASO monotherapy, 4 genes were expressed significantly different compared to the *mdx*-saline mice at FDR < 0.05. Two genes were unique to ASO monotherapy *Trdn* (−31%) with a role in RyR2 calcium channels and *Gm2a* (+76%) with roles including immune neutrophil function (Table 1). Two genes *Oasl2 and Irf7* were shared in the ASO+PMO combination and the MM+PMO combination groups (FDR < 0.05) and increased by similar levels in all 3 groups suggesting *Irf7* non-specific interferon effects via the phosphorothioate backbone in the oligonucleotides in these 3 groups (Table 1).

**Table 1.**
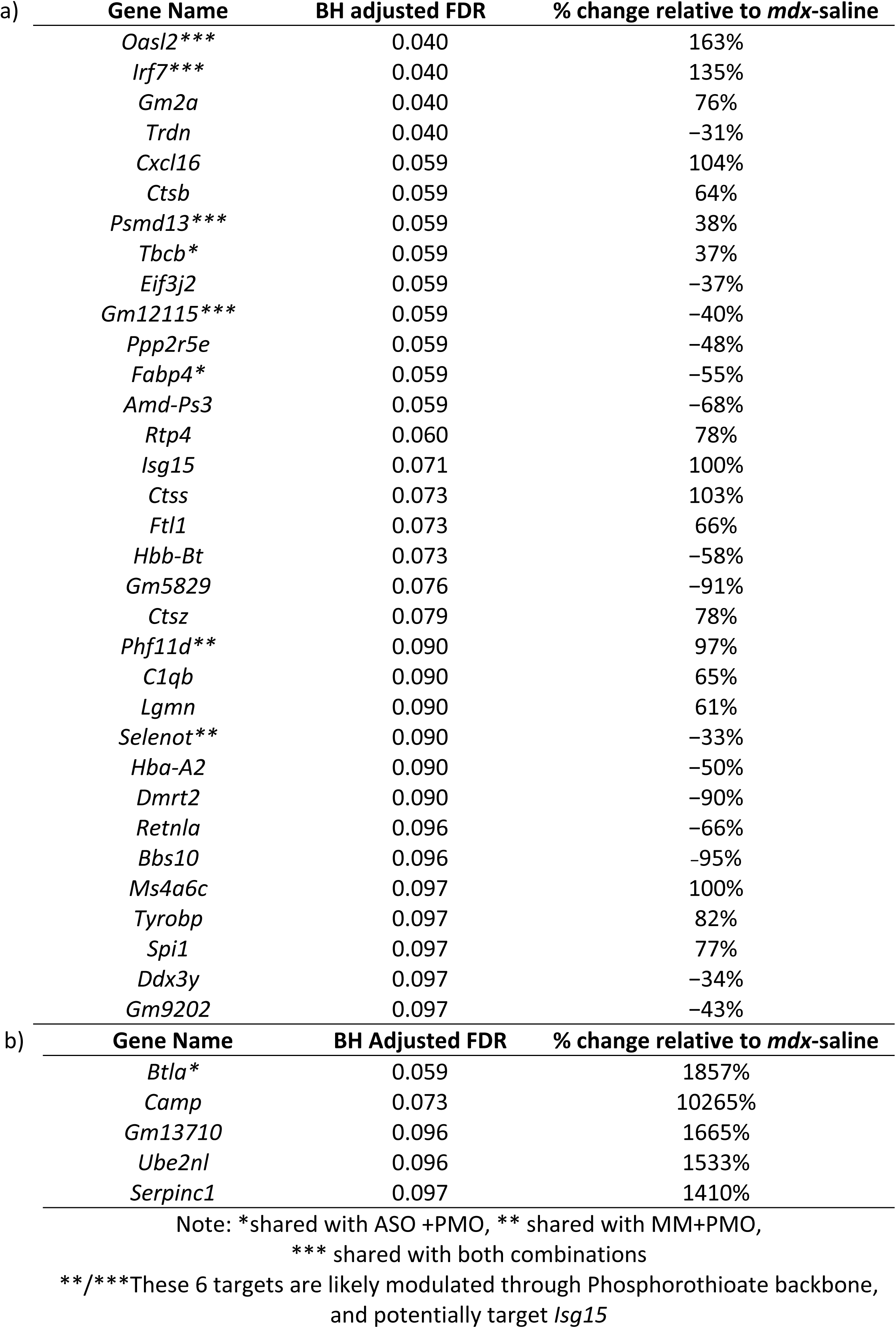
a) Summary data on genes with FDR < 0.1 in the ASO monotherapy group in order of the ascending FDR and b) Genes at the limit of detectability with RNA seq with FDR <0.1 in the ASO monotherapy group

In the ASO monotherapy at FDR < 0.1, a total of 38 targets were identified (Tables 1a and 1b), 29 of which are unique to the ASO monotherapy, 3 shared with the ASO+PMO combination (*Btla, Fabp4, Tbcb*) indicating relevance. Six were shared in groups with phosphorothioate backbone oligonucleotides; 4 of the 6 observed in both combinations (*Oasl2, Irf7, Psmd13, and Gm12115)*, and 2 of the 6 in the MM+PMO (*Phf11d, and Selenot*), suggesting non-specific modulation by the phosphorothioate in common with oligonucleotides in these 3 groups and for PHF11D potentially TLR-induced expression [41] (S2 Fig.).

Table 1a shows the 33 genes present starting at the lowest FDR, and Table 1b shows the additional 5 targets at the limit of detectability in *mdx* mice. The gene changes in Table 1a were in the range of +104% for *Cxcl16* to −95% for *Bbs10* for the genes specifically modulated by ASO versus saline in *mdx* mice. Gene changes in Table 1b were in the range of 14-fold to 102-fold, but these genes are expressed in the *mdx* saline samples at or below the limit of detection so the level of the fold change in the ASO group, may not be as accurate.

ASO monotherapy at FDR < 0.06 had an additional 10 genes versus FDR <0.05 (i.e. 14 in total) different from the *mdx*-saline mice. Of the 10 extra genes, 5 were unique to the ASO monotherapy, 2 of which were increased *Cxcl16* (+104%) a M2 chemokine involved in satellite myofiber regeneration and in *Cxcl16* knockout mice increased TGFb1 fibrosis [42], and *Cstb* (+64%) in neutrophil degranulation localized in the myoblast fusion stage of recovery with regenerating muscle fibres [43], with 3 decreased; *Eif3j2* (−32%) in initiation of protein synthesis, including of MHC Class II antigen processing [44], *Ppp2r5e* (−48%) suggesting glycogen metabolism to synthesize ATP and *Amd-PS4* (−68%) a pseudogene that does not produce protein.

Of the 10 extra genes, 3 in ASO monotherapy were shared with ASO+PMO combination namely, *Fabp4*, *Tbcb* and *Btla* (Table 1). Fatty acid binding protein 4, *Fabp4* reduced (−55%) in the monotherapy and (−49%) in the combination, is a gene expressed by the muscle adipose [45–47] modulated via CD49d+ M1 macrophages suggesting less fat in the ASO treated skeletal muscle [46,47]. *Tbcb* increased (+ 37%) in the monotherapy and similarly (+27%) in the combination, has a role in monocyte phagocytosis, and microtubule formation and stability [48]. *BTLA* an immune checkpoint inhibitor which suppresses B and T-cell activation and proliferation [49], had an 18.6-fold increase in the monotherapy (11.8-fold in the combination) with this level of increase less reliable as the gene was at the limit of detection in saline treated *mdx* mice.

The other 4 targets at the limit of detectability were unique to the ASO monotherapy and all increased versus *mdx* saline, and included Cathelicidin antimicrobial peptide (*Camp)*, that showed the largest 102-fold increase, with a role in innate immunity, chemotaxis and inflammatory response [50] increasing post-acute muscle injury via infiltrating activated neutrophil [51,52].

Low abundance target *Serpinc1* (antithrombin III) was increased 14-fold with a role in inhibition of thrombin activation of endothelia to interfere with VCAM-1 expression and CD49d+ monocyte adhesion [53,54] and IGF-1 transport [54]. *Igf-I* was upregulated 90% in *mdx* versus WT mice, as also observed in published studies.

Of the more abundant genes *Ddx3y* (−34%) is a Y linked gene expressed solely in male muscle reduced via interferon β1 expression during immune response [55]. *Tyrobp* (+82%) has a role in bone modelling as a receptor for Trem2, with a role in actin reorganization associated genes [56] associated with immune cell pathways in patients with DMD. Increased *Spi1* (+77%) reflected activated genes in the myeloid and lymphoid lineages such as CSF1R useful as a biomarker in DMD [56].

### Gene expression in the PMO monotherapy or MM monotherapy, and combination versus saline treated *mdx*

No significant differences in gene expression were observed in the *mdx* MM negative control oligonucleotide versus *mdx* saline at an FDR <0.1. Differences in gene expression were not observed in the PMO monotherapy versus *mdx* saline at an FDR < 0.1. In contrast, *Cacna1s*, was increased (46% and 53%) at an FDR <0.05 in the MM+PMO (FDR=0.025) and in the ASO+PMO (FDR=0.0079) combination respectively and was not seen in the ASO monotherapy nor MM monotherapy. This suggests *Cacna1s* modulation was related to the PMO in both combination samples. Similarly, genes *Gm8934* (−41%), and *Phldb1* (+54%) were modulated in the MM+PMO and the ASO+PMO combinations at FDR < 0.1 and not in the ASO nor MM, suggesting these changes were also related to the PMO in both combinations.

### ASO + PMO combination gene expression effects linked to immune function, lipolysis, fibrosis and muscle stem cells and muscle synthesis

The ASO+PMO combination at FDR < 0.05 showed 151 genes that were significantly different versus saline *mdx* mice, with 79 in ASO+PMO when compared to MM+PMO at FDR <0.05 (S1 Fig.), and 55 in the ASO+PMO combination at FDR < 0.05 when further compared to the MM+PMO combination up to FDR <0.1 (Table 2). Of these 55 differentially expressed genes 2, *Btla* and *Fabp4* are shared with the ASO group, at FDR at<0.06 and 53 are unique to the ASO+PMO combination at FDR <0.05.

**Table 2.** Summary of genes with FDR < 0.05 in the ASO+PMO combination unique from the MM+PMO at FDR<0.1 including the genes shared with ASO onotherapy at FDR <0.1 (*Btla,* and *Fabp4*).

| Gene name | BH adjusted FDR | % change relative to *mdx*-saline | Gene name | BH adjusted FDR | % change relative to *mdx*-saline | Gene name | BH adjusted FDR | % change relative to *mdx*-saline | Gene name | BH adjusted FDR | % change relative to *mdx*-saline |
| --- | --- | --- | --- | --- | --- | --- | --- | --- | --- | --- | --- |
| *Popdc3* | 0.0075 | −35% | *C230086J09Rik* | 0.028 | −94% | *Gm5268* | 0.036 | 272% | *Stbd1* | 0.042 | −39% |
| *Sh3rf2* | 0.0075 | 126% | *Etfrf1* | 0.029 | 58% | *Ap3s1-Ps1* | 0.036 | −94% | *Gm48909* | 0.047 | −32% |
| *Rpl11* | 0.013 | −32% | *Ppm1l* | 0.029 | 50% | *Gm48557* | 0.036 | −32% | *Btla* */** | 0.048 | 1184% |
| *Gm7908* | 0.015 | −67% | *Rbm25* | 0.029 | 30% | *Mt-Tp* | 0.036 | 114% | *Pimreg* | 0.048 | −92% |
| *Pik3ip1* | 0.018 | 91% | *Gm11425* | 0.03 | −39% | *Adam10* | 0.039 | −46% | *Gm10010* | 0.048 | −94% |
| *2310001k20rik* | 0.018 | 235% | *Fabp4* * | 0.030 | −49% | *Cbx3-Ps6* | 0.039 | 75% | *Gm6222* | 0.049 | −37% |
| *Gm5621* | 0.019 | −46% | *Gm29438* | 0.031 | −96% | *Gm4963* | 0.039 | −37% | *Mt-Atp6* | 0.049 | 57% |
| *Retreg1* | 0.02 | 54% | *Klf9* | 0.031 | 55% | *Gm5915* | 0.039 | −43% | *Gm37124* | 0.049 | −90% |
| *Calu* | 0.02 | −34% | *Rflnb* | 0.033 | −42% | *Supt5* | 0.039 | 36% | *Gm8894* | 0.049 | −62% |
| *Asb15* | 0.021 | 100% | *Igfbp7* | 0.034 | −34% | *Synpo2* | 0.039 | 39% | *Myom1* | 0.049 | 33% |
| *Gm43841* | 0.021 | 75% | *Klhl21* | 0.036 | 56% | *Tnfsfm13* | 0.039 | −91% | *Gm10123* | 0.049 | −41% |
| *Samd15* | 0.021 | −92% | *Lyz1* | 0.036 | −42% | *Gm4613* | 0.04 | −33% | *Gm7432* | 0.049 | −43% |
| *Rpl27-Ps1* | 0.023 | −29% | *Gm6012* | 0.036 | −90% | *Ubap2* | 0.041 | 46% | *G0s2* | 0.049 | −61% |
| *Sfr1* | 0.025 | 35% | *Gm8277* | 0.036 | 108% | *Mpc1* | 0.042 | 68% |  |  |  |

\* Two targets are shared with ASO monotherapy FDR <0.1. \*\*BTLA was at the limit of detectability in MDX saline mice.

Of 53 genes uniquely modulated by the ASO+PMO combination FDR < 0.05, it was noted that *Tnfsfm13* (−91%) and *Ppm1l* (+50%) play roles in macrophage activity and immune signaling [57,58]. *G0s2* (−61%) is a negative regulator of lipolysis in adipose tissue. *Igfbp-7* (−34%), is a fibrosis gene secreted from adipose-derived stem cells [59], and modulator of insulin-like growth factor (IGF) signaling pathway which increases muscle growth [60]. *Calu* (−34%) is associated with fibrosis and modulates the expression of IGF receptors and downstream signaling components.

*Asb15* (+100%), plays a role in muscle hypertrophy and is expressed predominantly in skeletal muscle [61]. *Myom1* (+33%), is essential in muscle development, stability, and function [62,63]. *Adam10* (−46%) is a regulator of satellite cell proliferation and differentiation, pharmacological inhibition of which enhances muscle regeneration after injury [64].

*Mt-Tp* (+114%) is involved in the incorporation of proline into muscles during development and maintenance with mutations associated with exercise induced fatigue and inflammation [65]. *Rflnb* (−42%) is a negative regulator of bone mineralization[66], suggesting potential for bone mineralization.

### ASO+PMO combination modulated genes at FDR <0.1 versus FDR <0.05

The ASO+PMO combination at FDR < 0.1 showed 203 unique genes that were significantly different expressed versus saline *mdx* mice, with only 3 shared with ASO group (*Btla, Fabp4, and Tbcb)* when compared to MM+PMO at FDR < 0.1. Thus, most of the ASO+PMO genes at FDR <0.1 are uniquely expressed compared to ASO. The extra 150 unique genes at FDR >0.05 to FDR <0.1 in the ASO+PMO, beyond those in Table 2, are in S1 Table. Pathway analysis done on the 203 unique targets is described below.

The ASO+PMO combination at FDR < 0.1 reduced *Cd163* FDR=0.068 (−61%) expressed by M2 macrophages expression in the combination, versus ASO monotherapy.

In the ASO+PMO combination at an FDR <0.05, there were 53 unique targets so few would be anticipated to be incorrect whilst between FDR >0.05 to FDR <0.1 there were an additional 150 genes, so potentially 15 incorrect genes which needs consideration when interpreting discoveries at the higher FDR.

### Pathway Analysis for ASO monotherapy and ASO+PMO combination gene expression effects demonstrated regulation of genes involved in immune and inflammatory pathways, lipolysis, fibrosis, and muscle cells

Pathway analysis was performed in Genetrails 3.2 on the 10 targets in the ASO FDR <0.06, 8 unique to the ASO therapy (*Gm2a, Trdn, Cxcl16, Cstb, Ptp2r5e, Eif3j2*, *Amd-Ps3* and *Tbcb*) including 2 (*Fabp4* and *Btla*) shared with ASO+PMO versus *mdx* saline (FDR< 0.05), related to ASO; *Tbcb* related to ASO is shared with the ASO+PMO at FDR<0.1 (S2 Table). Pathways identified included various immune and response to TNF pathways (*Cxcl16* and *Fabp4)*.

Pathway analysis using Genetrails 3.2 was performed on the 53 targets in the ASO+PMO combination FDR <0.05 versus saline (i.e. when excluding those in MM+PMO FDR < 0.1) (S3 Table) and on the 79 targets unique in the ASO and PMO combination FDR<0.05 (i.e. when excluding those in MM+PMO FDR < 0.05) with similar results (S1 Fig.). Pathway analysis for the 53 targets identified major regulation of Insulin-like Growth Factor (IGF) pathways, transport, and uptake as well as post-translational protein phosphorylation, and exercise induced circadian regulation.

The results using Genetrails3.2, and identified genes involved in immune, fibrosis, adipose and muscle stem cell and muscle cell pathways are represented in Fig. 4 [67].

**Fig. 4.**
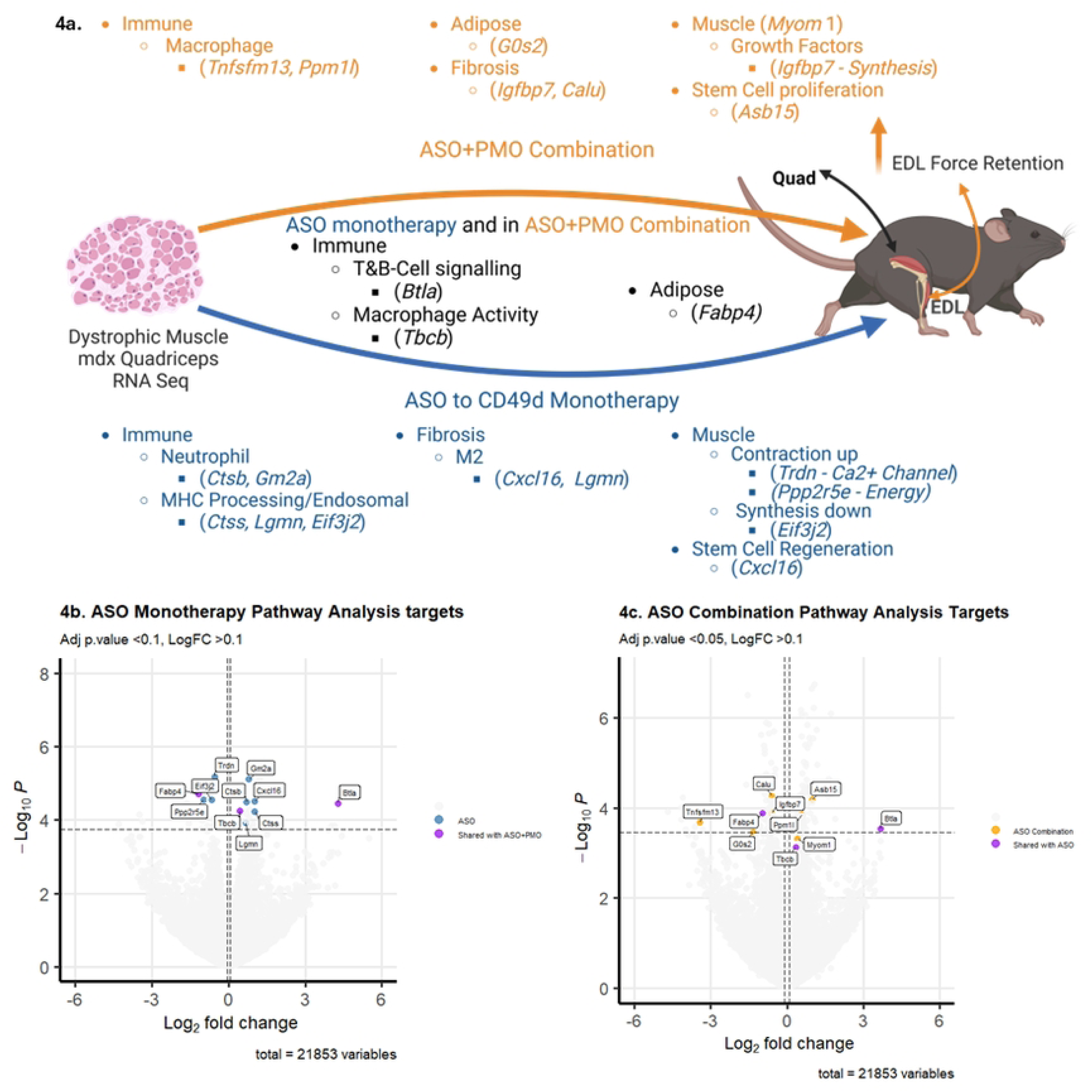
a) Summary of results from RNA seq and Pathway analysis using Genetrails 3.2 showing the genes involved in immune, fibrosis, adipose and muscle pathways modulated with the ASO onotherapy and ASO+PMO combination and the 3 shared (*Btla, Fabp4, Tbcb*). Volcano plot of the Genes affecting the pathways in 4a in the ASO monotherapy and c) Volcano ot of the Genes affecting the pathways in 4a in the Combination ASO+PMO and the 3 shared. e log2 FC indicates the mean expression level for each gene with the cut-off being LogFC > 0.1. ch coloured dot represents one gene, white dots represent genes not represented in the levant pathways. The Benjamini-Hochberg Adjusted p-value cutoff was b) p<0.1 and c) p<0.05.

ASO monotherapy modulated 8 genes referred to in Fig.4 of which 5 increased vs *mdx* saline, *Ctsb* (54%), *Gm2a* (76%), *Ctss* (103%), *Lgmn* (61%) and *Cxcl16* (104%) and 3 decreased, *Trdn* (-31%), *Ppp2r5a* (-48%), and *Eif3j2* (-37%). The 3 modulated by both ASO and the ASO+PMO combination showed similar level of modulation in both treatments *Fabp4* (-55%, -49%), *Tbcb* (37%, 27%) and *Btla* (18.6 fold, and 11.8 fold). ASO+PMO monotherapy modulated 7 genes referred to in Fig.4 of which 3 increased, *MyoM1* (33%), *Asb15* (100%), and *Ppm1l* (50%), and 4 decreased, *Tnfsfm13* (-91%), *G0s2* (-61%), *Calu* (-34%), and *Igfbp7* (-34%), the latter reduction potentially freeing increased IGF-I in mdx mice versus WT for muscle growth.

### ASO monotherapy modulated additional immune pathways and ASO+ PMO combination modulated additional immune, fibrosis, lipolysis, and muscle cells pathways

Pathway analysis using Genetrails3.2 was additionally performed on the 32 targets in the ASO monotherapy at FDR <0.1 (including the 3 shared with ASO+PMO combination at <FDR 0.1 (namely *Fabp4, Tbcb*, and *Btla*, as they are related to ASO) versus *mdx* saline. Pathways identified included various additional immune pathways in MHC class II antigen presentation, neutrophil degranulation, microglia pathogen phagocytosis pathway, Type II interferon signaling and complement, and coagulation cascades compared to results at FDR <0.05 (S4 Table).

Pathway analysis using Genetrails 3.2 was performed on the 203 targets unique in the ASO+PMO combination FDR <0.1 versus saline (when excluding MM+PMO FDR < 0.1) (S2 Fig.). Pathways identified included additional immunity pathways affecting TNF and inflammatory response, fibrosis, lipolysis, IGF-I, and muscle cell differentiation, regeneration, and development pathways compared to results at FDR <0.05 (S5 Table). The numerous ASO+PMO genes uniquely modulated by the combination suggest a synergy in the repair and regeneration of muscle.

## Discussion

This study is the first to investigate the effects of a unique combination treatment using a dystrophin morpholino exon skipping restoration drug (PMO) directed to the primary cause of damage in DMD together with an immune-modulating antisense drug (ASO) targeting adhesion molecule CD49d expressed on immune cells directed to secondary immune mediated damage in DMD. As several PMO drugs are conditionally approved for use by subsets of patients with DMD and ATL1102 to CD49d is showing promising stabilization effects in non-ambulant DMD patient trials, the potential effects of combining was investigated in *mdx* mice using the murine equivalents. The murine equivalent to ATL1102 to CD49d has previously been used as a monotherapy in *mdx*, and the murine PMO drug previously used in monotherapy in *mdx*, and this study sought to investigate effects at a PMO dose and at a time four weeks post dosing where low levels of dystrophin restoration occur, as observed in patients with DMD treated with known PMO restoration drugs. The ASO+PMO combination data supports the development of ATL1102 for the treatment of DMD in combination with conditionally approved PMO restoration drugs.

### Dosing with ASO to CD49d (ISIS348574), or PMO reduced AUC EC damage in EDL

In this study ASO monotherapy and the PMO monotherapy reduced EC damage in the EDL versus saline as assessed using the area under the curve of EDL function. The ASO to mouse CD49d dosed in symptomatic 9-week-old *mdx* mice once weekly for 8 weeks increased the area under the curve EDL function remaining after 10 eccentric contractions vs saline and MM control. The EDL was stretched *ex vivo* 10% each time with a short 3 minute recovery time in the 7-9 mice per group at ∼16 weeks of age. In a previous ASO monotherapy study in 9-week old mdx 12 mice per group were dosed once weekly for 6 weeks [17] EC recovery of the *tibialis anterior* was ∼25% higher versus saline after stretching the muscle *in situ* 5,10,15,20,25 and 30% with a 5 minute recovery. The differences in functional effect may be related to the different muscles tested, the *ex vivo* vs *in situ* studies, the timing of recovery post EC, and the age of the mice regeneration occurring at 8-12 weeks, at which time the greatest number of CD49d+ cells are observed in *mdx* versus WT week [69,70] whereas by week 16 there is fibrosis in the quadriceps [71].

The PMO in this study showed that 50mg/kg doses once weekly iv into the tail vein for 4 weeks, assessing four weeks past the last dose, improved area under the curve, EDL function remaining after 10 eccentric contractions in mice. Previous studies with this PMO show three once weekly higher 2mg iv injections in young 6 week old mdx mice (∼80mg/kg), 2 weeks after the last dose, leads to high 10-20% WT mice dystrophin levels in the *tibialis anterior* and provides higher maximum isometric tetanic force in this muscle 2 weeks past the last dose [19]. This PMO at the same 50mg/kg dose as used in the present study but in 12-month old mice, at 3 weeks after the last localized intra-arterial dose improves *ex vivo* EDL twitch tension mN/mm2 with average 8.6% dystrophin positive fibers in the EDL and 3.8% in the quadricep [18]. The present 50mg/kg PMO EDL study effects at 4 weeks post the last dose in ∼16 week old mice, thus extended previous observations at 2 weeks in 11 week old mice, and by 1 week vs 12 month old mice EDL functional data with this PMO in mdx, at a time when dystrophin is less than 1% WT in the quadriceps. A less than 1% WT dystrophin level in mdx is similar to dystrophin levels observed with eteplirsen and golodirsen in boys with DMD vs healthy control subjects and the present PMO mdx AUC EDL data suggests slight muscle functional protection in reduced eccentric muscle damage at such low dystrophin levels.

### Dosing with ASO+PMO combination improved maximum specific force and reduced damage after multiple EC in EDL

This *mdx* study demonstrated that once weekly dosing of symptomatic 9-week-old mice with a combination of ASO to CD49d for 8 weeks and PMO for the first 4 weeks improved at 8 weeks the specific maximum force and eccentric muscle force in the EDL muscle.

Comparatively, dosing 5-week old *Mdx* mice prednisolone at a human equivalent dose for 8 weeks did not reduce EDL eccentric muscle force loss but improved the specific maximum force [68].

In this study it was possible to see protective effect of ASO + PMO combination versus saline using EDL force remaining at 5 EC and AUC EDL function remaining after 10 EC. In the AUC EDL function analysis, the ASO+PMO combination produced statistically significant better effects than the PMO, trending better versus ASO monotherapy after 10 EC and statistically significantly better after 8 EC, the PMO producing significantly better effects versus the saline control and the ASO significantly better effects than both the saline and MM control in the area under the remaining EDL function curve.

No obvious changes in dystrophin protein production were observed in the *mdx* treated muscles with the PMO or PMO+ASO combination in the sample taken day 31 post PMO treatment. Of note this PMO, used at the same dose, produces a distinguishable ∼4% mean dystrophin expression compared to healthy WT dystrophin protein levels at day 17 post PMO treatment, and lower levels of less than ∼1.5% in healthy WT mice (Novak et al in supplemental Fig. 2.B) at day 31 [20], which suggests dystrophin restoration at earlier 2-3 week time points post PMO treatment also in this study.

This study shows that at a time when the ASO+PMO combination in *mdx* mice produces less than 1% WT dystrophin level in the quadriceps, it provides additional EDL muscle functional protection in maximum specific force and reduced eccentric muscle damage compared to saline and in the AUC vs monotherapy.

To explore the impact of ASO, PMO and ASO+PMO combination, further RNA sequencing gene expression analysis was conducted in the quadriceps of 16-week-old *mdx* mice versus WT mice and the multiple quadriceps gene expression benefits observed are described below in immune, fibrosis, and muscle cell and stem cell pathways.

### Transcriptomic analysis of WT versus *mdx* mice show alterations in calcium myonecrosis, inflammation, regeneration, fibrosis, adiposity, stem cell pathophysiology

The onset of disease in *mdx* mice is at 2 weeks and the peak of myonecrosis occurs at 4 weeks, with regeneration occurring at 8-12 weeks, at which time the greatest number of CD49d+ cells are observed in *mdx* versus WT [69,70]. By week 16 there is fibrosis in the quadriceps [71] and necrosis, and by week 20 necrosis is diminishing progressively with a little survival to week 26. in this study, there was substantial differentiation in the expression of fibrosis genes in 16-week-old *mdx* mice versus WT included VLA-4 ligands, *Spp1* (up 69-fold), *Fn1* (up 3.7-fold) and *Tgfβ1* (up 4.2-fold). Increased expression of genes in *mdx* versus WT mice including *Fn1* for fibrosis and *Spp1* for calcification are reported in areas of muscle regeneration [38]. Of note, fibrosis gene *Ctgf* was not different in the 16-week-old *mdx* vs WT mice and the CTGF targeted drug Pamrevlumab failed in phase III in ambulant and non-ambulant boys with DMD [39]. In contrast, the immune fibrosis associated gene *Hdac1* increased 8-fold in the *mdx* vs WT mice, and the HDAC1 targeted drug Ginivostat [40] is the only treatment to meet the primary endpoint in phase III ambulant boys DMD.

Dystrophic mice are characterized by calcium contraction induced muscle damage due to the loss of dystrophin, which leads to inflammation and further damage to muscle and the eventual replacement of muscle tissue with fibrotic and/or adipose tissue and the reduction in muscle stem cells [71]. Chronic muscle damage leads to cycles of activation of the innate immune system including neutrophils and macrophages and followed by persistent muscle antigen presentation, which leads to activation of the adaptive T and B cell immune system.

In this study, the ASO and the ASO+PMO combination modulated quadriceps genes in these pathological steps, but mostly via a different set of genes. Of 3 genes modulated by ASO in common to an FDR <0.1, 3 genes have roles in immune function, *Btla*, *Tbcb, Fabp4* and 1 gene also has a role in adipogenesis, *Fabp4*.

### ASO monotherapy modulated in the quadriceps immune, fibrosis, and muscles stem cell genes

The ASO to mouse CD49d monotherapy in the quadriceps uniquely modulated 2 genes at FDR<0.05. *Gm2a* was increased by 76% with a role including in immune neutrophil function and associated via binding to tetraspanin with activation of macrophages into anti-inflammatory M2 macrophages and their retention in tissues [72]. Mouse neutrophils express VLA-4 (CD49d, CD29) and degranulation involves CD29 [73]. The ATL1102 ASO to CD49d reduces CD49d RNA and surface expression of alpha-4 chain CD49d and beta-1 chain CD29 of VLA-4 [74] in line with the *Gm2a* effects in the study. GM2 inhibits growth factor receptor and cell adhesion, motility via integrin, by blocking them or their cross-talk, and this is enhanced in a complex with a tetraspanin, [75]. *GM2a* binding to tetraspanin (CD82) is required for muscle stem cell activation, and CD82 is decreased in human DMD [76]. *Trdn* was reduced 31% which can indicate open RyR2 calcium channel function [46,77] potentially previously closed via CD29 function, the beta chain of VLA-4 [78] which the ASO can modulate [74]. *Cacna1s*, a different calcium channel identified in the PMO+MM and PMO+ASO groups both at FDR <0.05 and not in any other groups, indicates its modulated by PMO in these combinations. *Cacna1s* is a calcium channel reported to work together with RyR1 calcium channel, so different calcium channel modulation featured with ASO versus PMO+ASO treatments [79].

The ASO monotherapy at FDR <0.1, modulated 29 unique genes in muscle versus the saline plus another 3 shared in ASO+PMO combination, 2 found in the ASO+PMO combination at FDR <0.05 (*Btla*, *Fabp4*) and 1 at FDR <0.1 (*Tbcb*). *BTLA* is a suppressor of B and T lymphocytes and increased >18-fold by ASO from a low abundance in *mdx*. *Fabp4* reduced 55% by ASO is expressed abundantly by adipocytes [45], modulated via CD49d positive M1 macrophage blockade suggesting ASO effects on M1 macrophages may reduce adiposity in muscle [47], with increased *TBCB* in muscle having a role in monocyte phagocytosis.

At FDR <0.06, with relatively low false discoveries (1 in 16 ), 7 targets were uniquely modified by the ASO, beyond *Trdn* and *Gm2a*. *Cstb* (+64%) involved in neutrophil degranulation, like *Gm2a*, is identified in *mdx* spatial transcriptomics in immune regions [38]. Cathepsin B, *(Ctsb)* peak is seen at the interface between inflammation and myoblast fusion [43]. *Ctsb* is a cysteine protease previously shown to be upregulated in DMD patient muscle, primarily localizing to macrophage infiltrating areas suggesting more of this myoblast fusion may occur in this *mdx* with ASO at week 16 [43].In contrast, mice treated with prednisolone for 3 weeks experience decreases in *Ctsb* and improved grip strength [43].

*Cxcl16* at FDR <0.06, an M2 macrophage chemokine with a role in muscle stem cell regeneration [78] was increased by 104% in the quadriceps compared to control and not modulated in the ASO+PMO combination, though different stem cell pathways are modulated in combination (Fig. 4). *Cxcl16* knockout mice have increased fibrosis linked to TGFβ1 expression [42] and *Cxcl16* triggers *b*1 integrin binding to VCAM-1. The CXCL16 receptor (CXCR6) expression is inversely correlated with another adhesion molecule CXCR4 [80] which increases post ATL1102 reduction of adhesion via CD49d and CD29 i.e. β1 integrin [74]. Notably ASO monotherapy increases of *Cxcl16* in *mdx* muscle are mirrored by increases in *CXCL16* in the plasma proteomics of adolescent boys with DMD treated with ATL1102 which stabilizes muscle function, strength, and reduces fat and increases lean muscle [81].

The ASO monotherapy at FDR <0.1 modulated 4 genes at the limit of detection in *mdx* versus the WT saline not modulated in the ASO+PMO combinations at FDR < 0.1. A known *mdx* low abundance gene *Camp*, showed the largest 102-fold increase, with a role in modulating innate immunity, chemotaxis, and inflammatory response. *Camp* increases occur post-acute muscle injury via infiltrating activated neutrophils [51] and activates classical macrophage expressing Mac-1 integrin β2and VLA-4 integrin β1 (CD29) binding to VCAM-1 [50]. Genetic knockout of *Camp* in mice ameliorates skeletal muscle pathology in *mdx* but *Camp* used as a treatment attenuates cardiac dysfunction in several preclinical models antihypertrophic effects protecting against *nfkappa* deleterious signalling [82]. *Camp* is described also as an agonist of IGF1R [83]. Another low abundance target *Serpinc1* (antithrombin III) increased 14-fold may indicate inhibition of thrombin activation of endothelia to interfere with VCAM-1 expression and CD49d+ monocyte adhesion [53,54]. *Serpinc1* additionally regulates IGF-1 transport and IGF 1 uptake by modulating IGF binding proteins [54]. *Igf-I* was upregulated 90% in *mdx* versus WT mice in this study, as observed in published studies, but beneficial effects are reported offset by *IGF-I* binding protein regulators [84]. Increases of *Serpinc1* in this study may enable ASO to CD49d to take advantage of increases in *Igf*-I.

ASO monotherapy at FDR <0.1 modulated *Ddx3y* (−34%) a Y linked gene expressed solely in male muscle and its reduction may reflect reduced interferon β1 expression during the immune response [55,84]. Interferon-β downregulates expression of *VLA*-4 and a reverse interplay may have occurred in this study [85]. ASO monotherapy also modulated *Tyrobp* (+82%) with a role in bone modelling, and in actin reorganization associated genes associated with immune cell pathways in patients with DMD, being one of 11 essential genes including *VCAM*-1 and *CSF1R* [86,87]. Increased *Spi1* (+77%) also occurred which activates genes in myeloid and lymphoid lineages such as *CSF1R* referred to as a Hub gene associated with pathology in boys with DMD useful as biomarkers [88].

Another gene notably modulated by ASO at FDR <0.1 was Legumain (*Lgmn*), an asparaginyl endopeptidase, a product of M2 macrophages with a role in antigen presentation, which increased by 61%. Legumain has a role in self-antigen presentation, and is associated with reducing collagen and fibronectin deposition [89]. This increase was mirrored in the plasma proteomics of ATL1102 in DMD patients with stabilized limb function and strength.

### ASO to mouse CD49d effects in *mdx* quadriceps parallels ATL1102 to human CD49d effects in DMD patients’ phase 2

ASO to mouse CD49d increased expression of *Cxcl16* (104%) and *Lgmn* (61%) in the *mdx* quadriceps have parallel changes in plasma of adolescent boys with DMD treated with ATL1102, with 28.9% higher CXCL16 and 42.2% higher LGMN in blood in the ATL1102 phase 2 study (data on file, human proteomics manuscript in preparation). CXCL16 is a M2 macrophage chemokine with a role in satellite cell myofiber regeneration [42] that triggers b1 integrin binding to VCAM-1 [74]. Legumain from M2 macrophages is associated with reducing collagen and fibronectin deposition [89].

There are potential parallels of reduced muscle fat content detected by MRI in the ATL1102 phase 2 with *Fabp4* reduced expression in the *mdx* by ASO to mouse CD49d. Increased *FABP4* is associated with increased fat accumulation in damaged muscle [90], and *FABP4* reduction suggestive of less fat and inflammation in skeletal muscle [91]. Osteopontin expression is linked to increased *FABP4* expression [47,92] and, in this study, osteopontin (*SPP1*) is 69X more highly expressed in *mdx* mice versus WT.

### ASO monotherapy shows a pattern of genes modulated in immune-inflammatory, fibrosis, calcium channel, and stem cell roles (different to those in ASO+PMO)

At FDR <0.1, 29 of 32 genes that were modulated in the ASO were the 2 neutrophil and M2 macrophage targets; *Ctsb* and *Gm2a* which were not modulated in combination. At FDR < 0.1, only 3 genes modulated by the ASO continued to be modulated by the ASO+PMO combination amongst 206 in total, 2 (*Tbcb* and *Fabp4)* potentially associated with M1 macrophage, *Fabp4* also expressed by adipocytes (Fig. 4). Thus, it appears a different muscle physiology exists in the ASO versus the ASO+PMO combination treated mice, with different pathways additionally affected in the combination.

Regulatory interactions between dystrophic muscle and the immune system involves neutrophils, followed by TH1 T cells and M1 macrophages involved in muscle repair, and then M2 macrophages in a wave [93]. All these pathways were affected with the ASO monotherapy. In the ASO monotherapy, neutrophil immune function *GM2a*, *CTSB*, together with macrophages function *TBCB*, both in the innate immune system, and T and B cell signaling in the adaptive immune system *BTLA*, were affected together with satellite cell *CXCL16* M2 regeneration and antifibrotic legumain M2 macrophage markers. *FABP4* in adipose potentially modulated by CD49d M1 macrophages and finally markers of improved muscle calcium *TRDN* channel and contraction *PPP2R5E,* suggesting more glycogen metabolism to synthesize ATP for contraction are observed, suggesting healthier muscle.

### ASO+PMO combination modulated an additional set of unique genes in the quadriceps muscle in immune cells, fibrosis, lipolysis, IGF-I, muscles stem cells and muscle Myomesin-1

The ASO+PMO at FDR < 0.05 modulated the most genes, 79 unique to the combination versus saline, not seen in MM+PMO at FDR < 0.05 (S1 Fig.). Of these, 55 are differentially expressed versus the MM+PMO at higher FDR <0.1 with 2 in common with ASO, *Btla*, and *Fapb4*. The 55 genes have roles in immune, fibrosis, lipolysis, IGF-1 and muscle stem cell physiology, and muscle growth (Fig. 4).

The ASO+PMO combination at FDR <0.1 generated 203 genes uniquely modulated versus saline (not seen in MM+PMO at FDR <0.1) with only 3 in common with ASO to FDR < 0.1, *Btla, the* T and B immune cells suppressor, *Tbcb* macrophage role, and *Fabp4* indicating lipolysis (S2 Fig.). The 203 genes modulated at FDR <0.1 had mostly similar physiological roles to roles identified for the 55 genes differentially expressed in the ASO+PMO at FDR < 0.05.

In the ASO+PMO *mdx* combination treatment at FDR <0.1 there were no neutrophil modulation genes, and the *Cd163* M2 marker gene was reduced (−63%), while it was not changed in the monotherapy and the combination had no M2 legumain or M2 *Cxcl16* changes found in the monotherapy consistent with reduction in *Cd163* in the combination.

In the ASO+PMO combination, there were additional macrophage TNF immune pathways, IGF-I for muscle protein synthesis and growth [94], and stem cell proliferation markers modulated when compared to the ASO monotherapy. These gene changes suggest muscle in a healthy state in the combination.

The ASO+PMO combination treatment resulted in a reduction in *Tnfsf13* (−91%) with the potential to reduce B cell and antibody producing plasma cells via receptor, *TACI*, and through the upregulation of NF-κB [57]. Increased *Ppm1l* implicated in regulating oxidative stress responses, prevents excessive inflammatory response in myocardial infarction through the inhibition of NF-κB [58]. Increased *Ppm1l* and reduced *Tnfsf13* suggests a potential reduced autoimmune response after dosing with combination treatment.

The ASO+PMO combination reduced *Igfbp7* known to be highly expressed by adipose derived stem-cells [59], high levels of *which are* associated with increased fibrosis and collagen content [95]. Additionally reduced *Igfbp7* can free IGF-I for improved muscle growth [60]. IGF-1 within muscle cells reduces necrosis of dystrophic myofibers in *mdx* mice and has the opposite effect to TNF which increases necrosis, with cross-talk between these molecules having implications in DMD [96].

The ASO+PMO combination increased *Asb15* by 100%, with a role in protein degradation within skeletal muscle cells. *Asb15* is part of the family of ankyrin repeat and SOCS box-containing proteins, that interact with proteins to regulate their stability or degradation. *Asb15* expression is upregulated during muscle regeneration, suggesting involvement in the repair and regeneration of damaged muscle tissue [61].

The ASO+PMO combination decreased *Adam10* otherwise capable of promoting satellite cells transition from a quiescent state to an activated state. A 48% decrease importantly favors satellite cells remaining in a quiescent state, maintaining the ability to replace progenitors. Suppression of *Adam10* activity in satellite cells can also activate satellite cells differentiation into myoblasts, which contribute to muscle repair and regeneration [64].

Elevated levels of *Adam10* have been seen in patients with idiopathic inflammatory myopathies, suspected to cleave FNDC5, which plays a key role in musculoskeletal metabolism [97].

The ASO+PMO combination increased *Myom1, the* Myomesin-1 protein involved in maintaining the structural integrity of muscle fibers [98]. *MYOM1* deficiency results in atrophic features and morphological abnormalities [99]. Increasing *MYOM1* expression, can enhance the stability and structural organization of muscle fibers leading to efficiency of force transmission improving muscle function in treated mice [63]. *MYOM1* has an integrin β1 talin binding site [100] the beta chain of VLA-4. *MYOM1* is increased by Ginivostat targeting HDAC1, recently registered after improving primary ambulation endpoints in DMD subjects [40,101] which PMO drugs have yet to achieve.

## Conclusions

The ASO to CD49d monotherapy effects include AUC EDL function changes and modulation of quadriceps gene expression of inflammatory and immune cells, fibrosis, stem cell markers, calcium channel and contractile muscle markers. The macrophage M2 related CXCL16, Legumain changes unique to the ASO monotherapy in the quadriceps have parallels with M2 changes seen in the plasma of ATL1102 treated non-ambulant DMD patients. It is possible such changes may have benefited the *mdx* mice *tibialis anterior* and non-ambulant DMD patients muscle function previously reported [17]. The ASO to CD49d plus PMO combination showed functional benefits in the *mdx* EDL and modulated 4 times the number of unique genes in the quadricep versus the ASO, many with known roles in improving muscle physiology. The pathways modulated uniquely in the ASO+PMO combination included additional immune response including B cell, fibrosis, lipolysis, IGF-I muscle growth factors, and muscle stem cell factors with increases in Myomesin-1 suggesting improved skeletal muscle physiology.

The upper limb function, strength and MRI muscle structure stabilization observed with ATL1102 monotherapy in phase 2 [17], and the *mdx* ASO+PMO combination functional EDL benefits and quadriceps differential gene expression pathway observations here-in support the investigation of ATL1102 in combination with the conditionally approved PMO drugs such as eteplirsen and golodirsen to improve the treatment outcomes of patients with DMD.

## Acknowledgments

The authors thank Susan Turner, Percheron Therapeutics Ltd, for logistical support in the acquisition of antibodies, oligonucleotides and the software for statistical work and thank IDT and Gene Tools for their expertise in the manufacture of the compounds used in this study. We also greatly thank Alli Patterson for reviewing and editing the manuscript for scientific content and clarity ahead of submission.

## Supporting Information

**S1 Fig. Two-set Venn diagram summarising at FDR <0.05 ASO+PMO combination gene changes.**

ASO+PMO had 79 unique gene changes, and another 72 changes shared with the MM+PMO groups. MM+PMO had 63 unique changes and of the 79 genes altered in the ASO +PMO combination 55 are unique versus MM+PMO FDR <0.1 and listed in Table 2.

**S2 Fig. Three set Venn diagram summarising at FDR <0.1 ASO+PMO combination gene changes**

ASO+PMO had 203 unique gene changes, 3* shared with ASO, 146 shared with MM +PMO combination groups, and another 4*** shared by all 3 groups related to PS. MM+PMO had 156 unique changes, and 2** shared between MM+PMO and ASO monotherapy also related to PS. The 38 genes modulated in the ASO monotherapy including the */**/***genes above are listed in Table 1.

## Notes

### Competing Interest Statement

I have read the journal's policy and the following authors of this manuscript have the following competing interests: Employed by Percheron Therapeutics: Padhye A S and Tachas G. Patent Application (inventors): Tachas G and Houweling P J.

